# The Omicron variant BQ.1* with mutations at positions 28,311 and 28,312 in the SARS-CoV-2 N gene have minimal impact on CDC N1 target detection

**DOI:** 10.1101/2023.01.26.525759

**Authors:** Guoping Ren, Bradley W. Langhorst, Gregory C. Patton

## Abstract

Ensuring COVID-19 testing remains accurate and reliable is of critical importance as the SARS-CoV-2 virus continues to evolve. Currently, a number of Omicron variants are dominating infection across the globe in including BQ.1 and XBB. Both variants and their sublineages (BQ.1* and XBB*) contain a 28,311 C/U mutation inherited from the original Omicron variant (BA.1). This mutation overlaps with a commonly used fluorescent probe for N gene detection in many Emergency Use Authorization (EUA) assays, as this target was originally established by the U.S. Centers for Disease Control and Prevention (CDC) in their EUA test for COVID-19 (2019-nCoV_N1). This C to U mutation was previously shown to have no impact on CDC N1 target detection. The rise of Omicron sublineages has increased the likelihood of additional point mutations occurring within the same assay target. A subpopulation of BQ.1* has an additional 28,312 C/U mutation within the CDC 2019_nCoV_N1 fluorescent probe in addition to the 28,311 C/U mutation. The double mutation could adversely affect the ability of diagnostic assays to detect the virus in patient samples and therefore it is important to verify the impacts of this additional mutation. Using in vitro transcribed (IVT) N gene RNA representing the wildtype (GenBank/GISAID ID MN908947.3) and Omicron BQ.1.1 variant (BQ.1, GISAID ID EPI_ISL_ 15155651), we evaluated the performance of two different amplification protocols, both of which include the CDC 2019-nCoV_N1 primer-probe set. Both assays successfully detected the mutant N gene sequence efficiently even at 10 copies of input, although the double mutation caused a 0.5∼1 C_q_ delay on average when compared to the wild-type sequence. These data suggest that circulating BQ.1* lineage viruses with this double mutation likely have minimal impact on diagnostic assays that use the 2019-nCoV-N1 primer-probe.

## Introduction

Ensuring that molecular diagnostic assays remain capable of detecting SARS-CoV-2 variants circulating in the patient population is critically important. New England Biolabs (NEB) previously established a workflow for monitoring and testing variants of SARS-CoV-2 that may impact commonly used primer and probe sets for diagnosis of COVID-19 (Figure 1). This workflow was used by NEB scientists to determine the impact of Omicron BA.1 on the CDC N1 target design, which has a C28,311U mutation in the genomic region targeted by the CDC N1 fluorescent probe^1^.

**Figure 1.**
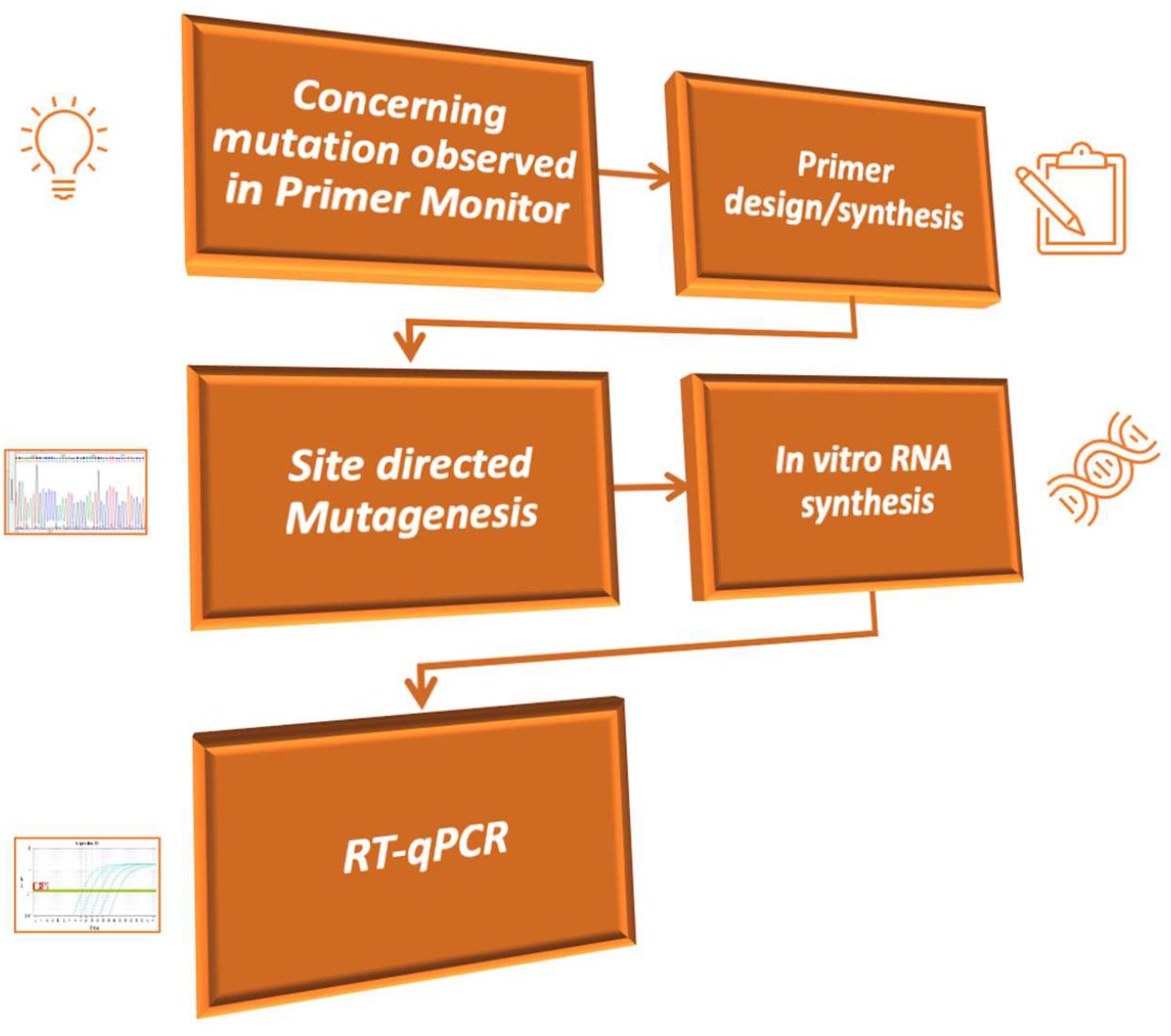
Monitoring variant impacts on RT-qPCR detection of SARS-CoV-2. Primer Monitor is a publicly available web tool that tracks SARS-CoV-2 virus variants and their impact on various diagnostic primer sets, including the CDC N1 & N2 targets. Once a concerning mutation(s) in the primer or probe assay design becomes predominate, its impact on the reaction can be determined by synthesizing mutant RNA *in vitro*. To this end, we designed primers to perform site-direct mutagenesis to generate the novel mutations in our N-gene encoded plasmid (N2117), which is used to create *in vitro* transcribed RNA. The purified RNA can then be used in an RT-qPCR assay with the impacted primers and/or probes to evaluate the effect of the mutation(s) on detection.

Recently, SARS-CoV-2 virus Omicron variant BQ.1 and its sublineages (BQ.1*) have been prevalent in global infections^2^. A small percentage of BQ.1* have a double mutation in the N1 probe region of the CDC assay design (C28311U, C28312U). We first noticed this variant in BQ.1* using our online Primer Monitor tool^3^. It highlighted a novel C to U mutation at position 28,312 in addition to the C to U mutation at position 28,311 from the BA.1 Omicron parental strain (Figure 2A). At the time of the writing, the frequency of this double mutation within sequenced SARS-CoV-2 samples globally over the last six months (07/18/22 to 01/17/23) is around 15%^4^. This newly introduced mutation is in the region of CDC 2019-nCoV_N1 probe target sequence. In combination with the previous 28,311 C/U mutation inherited from BA.1, this C to U double mutation (UU, hereafter) sits at the 3^rd^/4^th^ nucleotide from the 5’ end of the 2019-nCoV_N1 probe target sequence (Figure 2B). The UU mutation sufficiently raises concern of decreasing the effectiveness of COVID-19 diagnostic tests, since many tests utilize the CDC N1 assay design.

**Figure 2.**
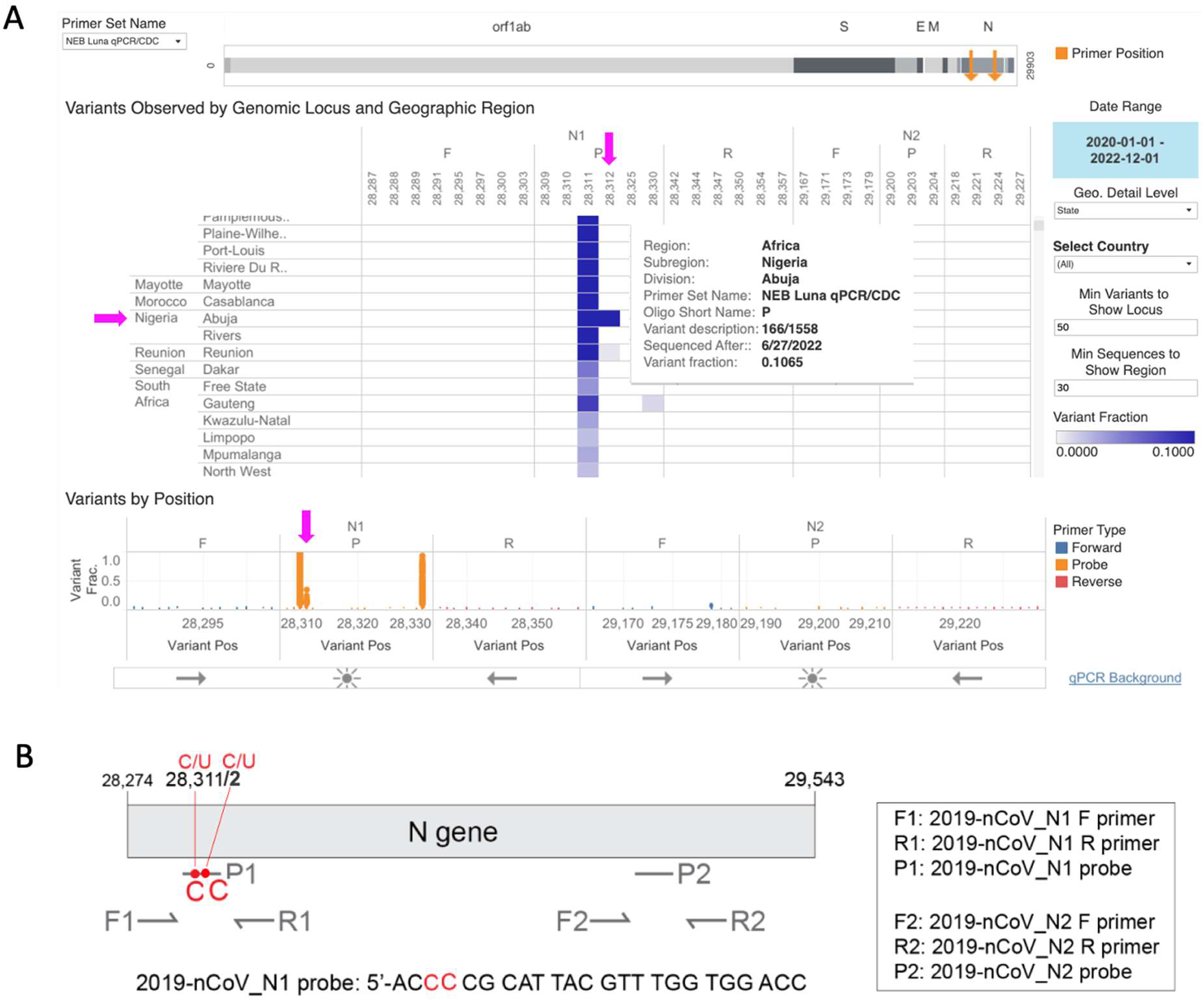
Primer Monitor Tool – BQ.1* variant contains UU mutation in the 2019-nCoV_N1 probe. A) Visual depiction of Omicron variant mutation within the N1 probe site generated using the Primer Monitor online tool (Tableau worksheet exported from Primer Monitor tool, primer set name “NEB Luna qPCR/CDC”). The UU double mutant was first noticed in SARS-CoV-2 sequences from Nigeria with a variant fraction around 10%. B) Schematic representation of the two CDC N gene primer probe sets. Each set includes one forward primer (F), one reverse primer (R), and one fluorescent probe (P). The SARS-CoV-2 BQ.1* variant has a C to U mutation at position 28,311 and 28,312, which is within the 2019-nCoV_N1 probe (P1) target sequence. Not drawn to scale.

Herein, we followed the previous workflow established to compare qPCR amplification efficacy for the CDC N1 target in wild-type and BQ.1 variant N gene synthetic RNA sequences using two different amplification protocols. We find a small C_q_ delay is observative for the double mutation, but the target is still detectable at 10 cp per reaction.

## Results

### NEB SARS-CoV-2 Multiplex Assay detects the N1 target from N gene RNA containing the BQ.1* variant mutation

The UU mutation at position 28,311/28,312 is likely to impact the N1 fluorescent probe annealing but required assessment to determine the overall effect on assays that rely on the CDC N1 design. In order to test the assay efficiency on this variant RNA, wild-type N gene and mutant RNA templates were made by IVT. Correct size and high purity were visualized via agarose gel (data not shown). We then compared amplification of the mutant RNA to the wild-type N gene RNA using the NEB SARS-CoV-2 RT-qPCR Multiplex Assay Kit (E3019), which uses the CDC N1 and N2 assay design for SARS-CoV-2 detection. This assay kit relies on multiplexing, simultaneously detecting N1 (HEX), N2 (FAM) and human RNase P (Cy5) in a single reaction. The N1 (Figure 3A) and N2 (Figure 3B) targets were amplified efficiently across a 7-log dilution (10^7^-10 copies/reaction) of the mutant and wild-type input RNA. In these experiments, the N2 primer-probe set also served as an internal control to monitor minor differences in RNA template input as the target is identical for both templates.

**Figure 3.**
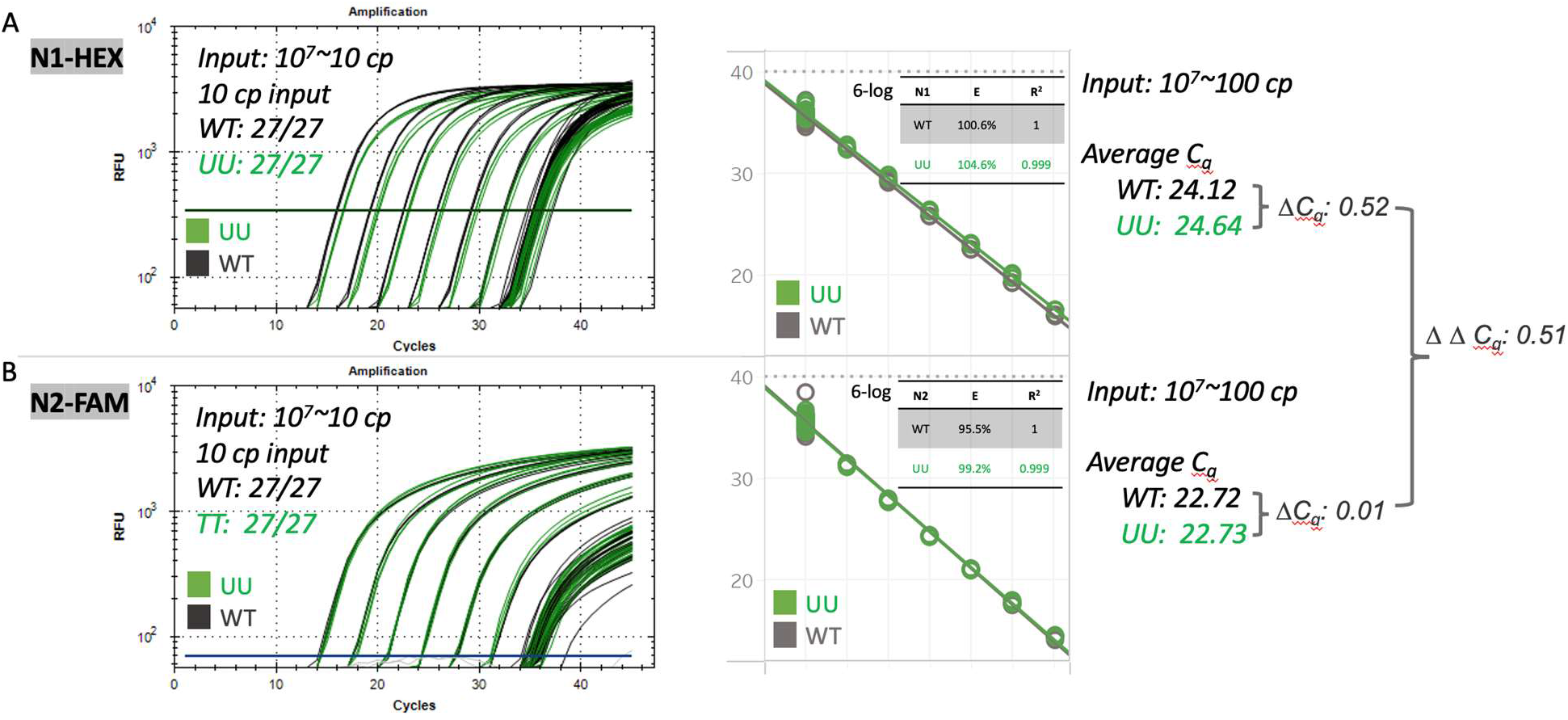
The CDC 2019-nCoV_N1 primer-probe set remains efficient in BQ.1* detection in the NEB SARS-CoV-2 Multiplex Assay. Amplification efficiency was evaluated in triplicate over a 7-log range (10^7^-10 copies/reaction) of synthetic wild-type RNA (black) versus the UU mutant RNA (green). Detection sensitivity was evaluated with 10 copies of RNA per reaction with 27 replicates per condition. NEB SARS-CoV-2 RT-qPCR Multiplex Assay Kit (NEB E3019) was used to detect A) N1 (HEX) and B) N2 (FAM) targets simultaneously following cycling conditions as recommended by the manufacture. All twenty-seven low copy replicates (10 copies) were detected successfully using CDC 2019-nCo_N1 primers and probe.

The C_q_ values for detection of WT and mutant RNA for N2 (FAM channel) were nearly identical (average ΔC_q_ = 0.01), ensuring experiments contained an equal amount of template RNA. N1 target amplification between the mutant and wild-type RNA showed small differences in C_q_ values, with average ΔC_q_ of 0.52 due to the higher C_q_ value for the mutant RNA (Figure 3A).

The ΔΔC_q_ is calculated as 0.51, reflecting a minor C_q_ delay for the mutant. The slower mutant RNA detection is likely due to slightly attenuated probe annealing related to the double mutation. More relevantly, we assessed the assay sensitivity by testing with low input RNA using 10 copies per reaction. Both the wild-type and mutant RNA were successfully detected by the N1 and N2 target designs, with 27 out of 27 replicates detected. These data suggest the NEB SARS-CoV-2 RT-qPCR Assay can still efficiently detect the IVT BQ.1 N gene double mutant using the CDC 2019-nCoV_N1 primer-probe set, albeit with a small C_q_ delay.

### N gene RNA containing the BQ.1 variant mutation can be detected using SalivaDirect™ RT-qPCR amplification conditions

SalivaDirect is currently a widely deployed real-time RT-qPCR SARS-CoV-2 diagnostic platform that allows for simple diagnosis of COVID-19 using saliva, rather than nasal swabs, as a patient sample^5^. It is a two-plex assay that has been granted EUA by the Food and Drug Administration (FDA) that leverages the CDC 2019-nCoV_N1 target and human RNase P as an internal control. Following the RT-qPCR protocol as outlined in the SalivaDirect assay, we observed efficient amplification of both the mutant and wild-type RNA across a 7-log dilution series of RNA using either Luna^®^ Probe One-Step RT-qPCR 4X Mix with UDG (NEB M3019, Figure 4A) or the Luna Universal Probe One-Step RT-qPCR Kit (NEB E3006, Figure 4B). The mutant RNA was detected slightly slower with an average ΔC_q_ of 0.87. However, this assay remains highly sensitive for the mutant N gene, as 10 copies of input RNA were detected in 27 out of 27 replicates tested. This is in consistent with the NEB SARS-CoV-2 Multiplex Assay data. We therefore conclude that both the NEB SARS-CoV-2 Multiplex Assay Kit and the SalivaDirect amplification conditions can reliably detect the BQ.1 N gene containing this double mutation using the CDC 2019-nCoV_N1 primer-probe set.

**Figure 4.**
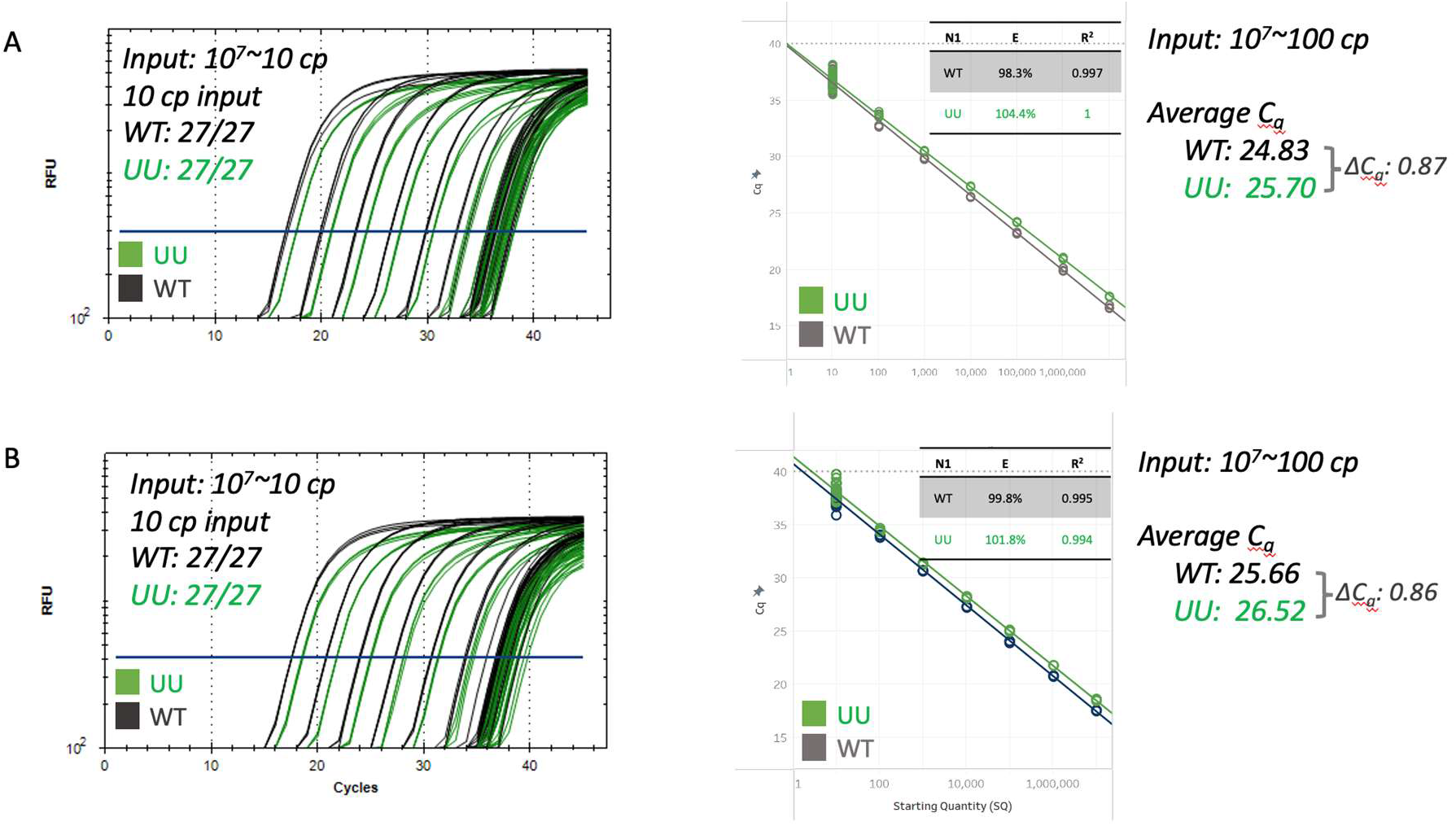
The CDC 2019-nCoV_N1 primer-probe set remains efficient in BQ.1* detection in the SalivaDirect RT-qPCR protocol. Amplification efficiency was evaluated in triplicate over a 7-log range (10^7^-10 copies/reaction) of synthetic wild-type RNA (black) versus the UU mutant RNA (green). Detection sensitivity was evaluated with 10 copies of RNA per reaction with 27 replicates per condition. A) Luna Probe One-Step RT-qPCR 4X Mix with UDG (NEB M3019) or B) the Luna Universal Probe One-Step RT-qPCR Kit (NEB E3006) were used to detect the N1 (FAM) and the RP (Cy5) targets simultaneously following the SalivaDirect RT-qPCR protocol. All twenty-seven low copy replicates (10 copies) were detected successfully using CDC 2019-nCo_N1 primers and probe.

## Discussion

New SARS-CoV-2 variants based on the original Omicron sequence (BA.1) continue to emerge across the world. Tracking the emergence of new variant mutations and their impacts to molecular assays will help ensure the continued reliability of COVID-19 diagnostic tests. To support this effort, NEB previously established a workflow to closely monitor the potential impacts of the mutations towards various molecular diagnostics and facilitate the evaluation process. The Primer Monitor tool (https://primermonitor.neb.com/) plays an extremely important role in automatically monitoring the mutational spectrum across a wide range of publicly submitted amplification primer sets. It has proved highly successful in calling out concerning mutations, including revealing the original mutation in Omicron (BA.1) at position 28,311, which overlapped with the CDC 2019-nCoV_N1 target sequence for detection of SARS-CoV-2 in real-time RT-PCR^6^. Some mutations have been found to decrease assay sensitivity (*e*.*g*., CDC N2 on G29179U) while others have had no observable impact on sensitivity (*e*.*g*., CDC N1 on C28311U or C28311U/A28330G)^1,3,6,7^.

The BQ.1* lineage with the UU double mutation is particularly concerning because it overlaps with the CDC N1 probe region, potentially decreasing the efficacy of COVID-19 diagnostic tests that employ this target design. Our performance evaluation of two different amplification conditions found that the N1 site is efficiently detected with the CDC 2019-nCoV_N1 primer-probe set, although the double mutation results in a small C_q_ delay. While additional testing is needed to determine if other workflows may be impacted, our results on purified RNA suggest robust amplification performance is maintained on the BQ.1 N-gene C to U 28,311/28,312 double mutant.

## Methods

The SARS-CoV-2 N gene RNA harboring the C to U mutation at position 28,311 and 28,312, was generated using site-directed mutagenesis (Q5 Site-Directed Mutagenesis Kit, NEB E0554S) of the SARS-CoV-2 Positive Control (N gene) plasmid (NEB N2117). Sanger sequencing confirmed successful mutagenesis. Wild-type or mutant plasmid was linearized using restriction digestion and then subsequently used for *in vitro* transcription (HiScribe^®^ T7 High Yield Synthesis Kit, NEB E2040) to synthesize wild-type and mutant N gene RNA, respectively. A Monarch^®^ RNA Cleanup Kit (NEB, E2050) was used to purify the RNA and the RNA was quantitated with the Nanodrop to calculate the RNA copy number. The purity and quality of the plasmids and RNA were assessed using agarose gel electrophoresis (1.2 %, 1 h, 0.5x Tris-Borate-EDTA Buffer).

The impact of the 28,311/28,312 C to U mutation on the CDC 2019-nCov_N1 target detection was conducted as previously described^6^. Briefly, a 7-log dilution series (10^7^-10 copies/reaction) was prepared for both the wild-type and the mutant RNA, with 10 ng of Jurkat total RNA (BioChain, R1255815-50) included as an internal control. The mutant and wild-type target RNA were subsequently amplified using either the NEB Luna SARS-CoV-2 RT-qPCR Multiplex Assay Kit (NEB E3019), Luna Probe One-Step RT-qPCR 4X Mix with UDG (NEB M3019) or the Luna Universal Probe One-Step RT-qPCR Kit (NEB E3006) following the standard NEB protocol or SalivaDirect RT-qPCR amplification protocol as indicated. For sensitivity evaluations, 27 reactions containing 10 copies/reaction were performed using either wild-type or mutant RNA. For the Luna SARS-CoV-2 RT-qPCR multiplex assay, the N1 (HEX), N2 (FAM) and RP (Cy5) targets were simultaneously detected using the following cycling conditions: carryover prevention (25°C for 30 s), cDNA synthesis (55°C for 10 min), initial denaturation (95°C for 1 min) and 45 cycles of denaturation (95°C for 10 sec) and annealing/elongation (60°C for 30 sec) plus a plate read step. For the SalivaDirect RT-qPCR, both the Luna Probe One-Step RT-qPCR 4X Mix with UDG (NEB M3019) and the Luna Universal Probe One-Step RT-qPCR Kit (NEB E3006) were used to detect the N1 (FAM) and the RP (Cy5) targets simultaneously using the following cycling conditions: cDNA synthesis step (52°C for 10 min), initial denaturation (95°C for 2 min) and 45 cycles of denaturation (95°C for 10 sec) and annealing/elongation (55°C for 30 sec) plus a plate read step. The qPCR data was collected on a Bio-Rad CFX96 qPCR instrument (96-well, 20 μl reactions).

## Acknowledgements

We thank the NEB DNA Sequencing Core for Sanger sequencing.

